# Th1 cells express Tfh effector molecules and associate with B cells at the T-B border following viral infection

**DOI:** 10.1101/2025.04.02.646697

**Authors:** Jennifer S. Chen, Abigail L. Tierney, Oluwagbemiga A. Ojo, Ryan D. Chow, Eric Song, Tianyang Mao, Benjamin Israelow, Jason S. Weinstein, Adam Williams, Craig B. Wilen, Stephanie C. Eisenbarth

**Affiliations:** Department of Dermatology, University of Pennsylvania; Philadelphia, PA, USA; Department of Medicine, Division of Allergy and Immunology, Northwestern University Feinberg School of Medicine; Chicago, IL, USA; Center for Human Immunobiology, Northwestern University Feinberg School of Medicine; Chicago, IL, USA; Department of Medicine, University of Pennsylvania; Philadelphia, PA, USA; Department of Ophthalmology, Yale University School of Medicine; New Haven, CT, USA; Broad Institute of MIT and Harvard, Cambridge, MA, USA; Department of Internal Medicine, Section of Infectious Diseases, Yale University School of Medicine; New Haven, CT, USA; Center for Immunity and Inflammation, Rutgers New Jersey Medical School; Newark, NJ, USA; Department of Immunobiology, Yale University School of Medicine; New Haven, CT, USA; Department of Laboratory Medicine, Yale University School of Medicine; New Haven, CT, USA

## Abstract

Antibodies protect the host from pathogens by neutralizing viruses, targeting pathogens for phagocytosis and complement deposition, and by activating innate immune cells. Most studies on the cellular mechanisms of antibody induction focus on the role of germinal centers (GCs) – areas within lymph nodes (LNs) and spleen where activated B cells undergo mutation and then selection by T follicular helper (Tfh) cells to produce high affinity antibodies. However, specific and long-lived antibodies can be produced in the absence of GCs and Tfh cells, sometimes referred to as “extrafollicular” (EF) responses. However, EF responses are typically assumed to be short-lived and low affinity. We characterized the CD4^+^ T cells present in wild type and Tfh cell-deficient mice in lung draining LNs following influenza A virus or SARS-CoV-2 infection to compare the nature of T cell help for activated B cells. We found that a population of helper Th1 cells remains in LNs seven days following viral infection and interacts with IgG2c^+^ B cells at the T-B border in both WT and Tfh-deficient mice. We propose that Th1 cells promote virus-specific antibody production in parallel with Tfh cells in the GC and that such a response is contiguous with the T-B border and therefore is not an EF response as classically defined.

## Introduction

### T cell dependent, high affinity antibodies to immune stimuli are induced canonically through T follicular helper (Tfh) cell and germinal center B (GCB) cell interactions

Tfh cells are uniquely equipped to aid and direct germinal center responses given their ability to enter the B cell follicle and directly interact with GC B cells. In the germinal center, Tfh cell help is limiting and is essential for driving selection of GC B cells. Classically, Tfh cells provide help to B cells in the form of CD40L and two key Tfh-associated cytokines, IL-4 and IL-21 (1). The combination of costimulation and cytokines promotes survival and the GC B cell program (i.e. BCL6 expression). B cells that have received Tfh help will continue to cycle through the germinal center, where they undergo iterative rounds of somatic hypermutation and proliferation until they exit the germinal center with highly mutated BCRs with improved ability to recognize and bind antigen. Upon GC exit, the B cells can either remain as memory cells or become plasma cells that secrete high-affinity antibodies that protect against pathogens.

### Extrafollicular B cell responses induce early, protective antibodies

A parallel B cell response occurring outside the GC, and termed extrafollicular (EF), is induced early prior to the formation of GCs and outputs protective memory B cells and antibody secreting plasma cells during a T-dependent stimuli (2–5). While the originally described EF B cell response was of rapidly dividing plasma blasts (PB) that provided an early antibody response in the red pulp, the EF term usage has expanded to include responses localized to regions bridging the B cell follicle in the spleen and medullary cords in the lymph node (LN) (2, 3, 6–10). In contrast to the prevailing view of a transient EF reaction during T-dependent responses, recent studies indicate GC-independent memory B cells are generated throughout an immune response (5). Due to the absence of Tfh-mediated selection, as present in the GC reaction, EF responses are classically thought to generate cells with low affinities. However, data from us and others clearly show class switched and somatically mutated cells with high affinities can be generated independently of Tfh cells during these responses (5, 11, 12).

### Genetically impairing GCs only impairs certain types of antibodies

In the absence of Tfh cells, following various models of viral infection in the lung or food allergen immunization through the gut, antigen-specific IgG1 and IgE antibodies are lost, while antigen-specific IgG2b, IgG2c and IgA remain intact (11–13). This implies that another cell subset, presumably a CD4^+^ T cell subset, is sufficient to provide the proper signals to induce class switching and selection in vivo. Indeed, studies have demonstrated that CD4^+^ T cell help is required for the generation of virus-specific IgG2 antibodies (11), and more specifically, IFN-γ-producing Th1 cells are sufficient to support IgG2 antibody production in the absence of Tfh cells and the GC (13, 14). Moreover, these Th1 cells have been shown to express the cytokine IL-21 and key costimulatory molecules such as CD40L and ICOS, all of which are classically associated with Tfh cells and their help of B cells during GC reactions (13). This implies that in the absence of Tfh cells, Th1 cells can compensate and provide B cells with the necessary signals for activation and class switching in a GC-independent manner.

### GC-independent cellular interactions can promote antibodies that are long lived and capable of neutralizing viruses

In one study, vaccination followed by lethal challenge with influenza A virus demonstrated that IgG2 antibodies produced in the absence of Tfh cells and the GC are sufficiently protective against influenza virus infection, although these antibodies were of lower avidity than antibodies made when Tfh cells were present. Additionally, these antibodies from Tfh-deficient mice had equivalent neutralizing capacity in vitro when compared to that of Tfh-sufficient mice (13). Similarly, in SARS-CoV-2 infection, Tfh-deficient mice generate high-affinity Spike-specific IgG antibodies that are detectable up to 3 months post infection with equivalent, if not improved, neutralization potency when compared to Tfh-sufficient mice (11). These data seem to challenge the notation that EF responses result in short-term, lower-quality antibody responses in the context of different viral infections.

We report here that BCL6-indepdent CD4^+^ T cells can provide Tfh-like signals capable of promoting B cell activation during SARS-CoV-2 and PR8 influenza infection. We observed a population of mixed Th1 cells that expressed CD40L and produced the cytokines IFN-γ and IL-21 in the presence or absence of Tfh cells. When Tfh cells were absent, this mixed Th1 population became the dominant expressor of CD40L and main cytokine producer in the lung draining LN. During SARS-CoV-2 infection, these CD4+ T cells associated with B cells outside of the GC in the lung draining LN regardless of whether Tfh cells and GCs were present or absent. Additionally, these T-bet^+^ non-Tfh cells were observed interacting with IgG2c^+^ B cells at the T:B border both in WT and Tfh-deficient mice. Taken together, these data imply that there are parallel pathways of antibody production occurring during viral infection, one that is defined as the classic GC/Tfh-dependent pathway and a second that is characterized by GC-independent/LN-resident Th1-mediated help of B cells. Ultimately, both pathways are sufficient to provide the host with long-lived, high-affinity, and protective antibodies during acute respiratory viral infection.

## Materials and Methods

### Mice

*Bcl6*^fl/fl^ [B6.129S(FVB)-*Bcl6*^*tm1*.*1Dent*^/J (15)] and *Cd4*^Cre^ [B6.Cg-Tg(Cd4-cre)1Cwi/BfluJ (16)] mice were purchased from Jackson Laboratory. *Bcl6*^fl/fl^ mice were crossed with *Cd4*^Cre^ mice to generate *Bcl6*^fl/fl^*Cd4*^Cre^ mice. Mice were bred in-house using mating trios to enable utilization of littermates for experiments. Mice of both sexes between 6 and 10 weeks old were used for this study. Animal use and care was approved in agreement with the Yale Animal Resource Center and Institutional Animal Care and Use Committee according to the standards set by the Animal Welfare Act.

### Cell lines

Huh7.5 and Vero-E6 cells were from ATCC. Cell lines were cultured in Dulbecco’s Modified Eagle Medium (DMEM) with 10% heat-inactivated fetal bovine serum (FBS) and 1% Penicillin/Streptomycin. All cell lines tested negative for *Mycoplasma* spp.

### AAV-hACE2 transduction

Adeno-associated virus 9 expressing hACE2 from a CMV promoter (AAV-hACE2) was purchased from Vector Biolabs (SKU AAV-200183). Mice were anesthetized by intraperitoneal injection of ketamine (100 mg/kg) and xylazine (10 mg/kg). The rostral neck was shaved and disinfected with povidone-iodine. After a 5-mm incision was made, the salivary glands were retracted and the trachea visualized. Using a 31G insulin syringe, 10^11^ genomic copies of AAV-hACE2 in 50 μl PBS were injected into the trachea. The incision was closed with 3M Vetbond tissue adhesive. Following subcutaneous administration of analgesic consisting of meloxicam (5 mg/kg) and buprenorphine XR (3.25 mg/kg), mice were placed in a heated cage and monitored until full recovery.

### Viruses

SARS-CoV-2 P1 stock was generated by inoculating Huh7.5 cells with SARS-CoV-2 isolate USA-WA1/2020 (BEI Resources, NR-52281) at a MOI of 0.01 for three days. The P1 stock was then used to inoculate Vero-E6 cells, and the supernatant was harvested after three days at 50% cytopathic effect. The supernatant was clarified by centrifugation (450 × *g* for 5 min), filtered through a 0.45-micron filter, and stored in aliquots at -80°C. For infection of AAV-hACE2 mice, virus was concentrated by mixing one volume of cold 4X PEG-it Virus Precipitation Solution (40% wt/vol PEG-8000 and 1.2 M NaCl) with three volumes of viral supernatant. The mixture was incubated overnight at 4°C and then centrifuged at 1500 × *g* for 60 min at 4°C. The pelleted virus was resuspended in PBS and stored in aliquots at -80°C. Virus titer was determined by plaque assay using Vero-E6 cells (17).

Influenza virus A/PR/8/34 H1N1 (PR8) expressing the ovalbumin OT-II peptide was grown for 2.5 days at 37°C in the allantoic cavities of 10-day-old specific-pathogen-free fertilized chicken eggs. Harvested virus was centrifuged at 3000 × *g* for 20 min at 4°C to remove debris and stored in aliquots at -80°C. Virus titer was determined by plaque assay on Madin-Darby canine kidney cells (18).

For all infections, mice were anesthetized using 30% vol/vol isoflurane diluted in propylene glycol (30% isoflurane) and administered SARS-CoV-2 or PR8 intranasally in 50 μl PBS. AAV-hACE2 *Bcl6*^fl/fl^ and *Bcl6*^fl/fl^*Cd4*^Cre^ mice were infected with 1.2×10^6^ PFU of SARS-CoV-2. *Bcl6*^fl/fl^ and *Bcl6*^fl/fl^*Cd4*^Cre^ mice were infected with 70 PFU of PR8. All work with SARS-CoV-2 was performed in a biosafety level 3 (BSL3) facility with approval from the office of Environmental Health and Safety and the Institutional Animal Care and Use Committee at Yale University.

### Flow cytometry

Mediastinal lymph nodes (medLN) were homogenized with a syringe plunger and filtered through a 70-μm cell strainer. Red blood cells were lysed with RBC Lysis Buffer (BioLegend) for 1 min. Single-cell preparations were resuspended in PBS with 2% FBS and 1 mM EDTA and pre-incubated with Fc block (clone 2.4G2) for 5 min at room temperature before staining. Cells were stained with the following antibodies or viability dye for 30 min at 4°C: anti-CD4 (RM4-5), TCRβ (H57-597), PD-1 (RMP1-30), CD44 (IM7), PSGL-1 (2PH1), Ly6C (HK1.4), B220 (RA3-6B2), Fas (Jo2), GL7 (GL7), CD138 (281-2), and LIVE/DEAD™ Fixable Aqua (Thermo Fisher). CXCR5 (L138D7) was stained for 25 min at room temperature.

CD40L staining was performed by surface mobilization assay (19). Cells were pre-incubated with Fc block (10 μg/ml) for 10 min at room temperature. Cells were then stimulated with 50 ng/ml PMA and 1 μg/ml ionomycin in complete RPMI (10% heat-inactivated FBS, 1% Penicillin/Streptomycin, 2 mM L-glutamine, 1mM sodium pyruvate, 10 mM HEPES, 55 μM 2-mercaptoethanol) in the presence of PE/Cy7 anti-CD40L (MR1, 4 μg/ml) for 30 min at 37°C. After washing, the remaining markers were stained as described above.

For intracellular cytokine staining, cells were stimulated with 50 ng/ml PMA and 500 ng/ml ionomycin in complete IMDM (10% heat-inactivated FBS, 1% Penicillin/Streptomycin, 2 mM L-glutamine, 1mM sodium pyruvate, 25 mM HEPES, 1X MEM Non-Essential Amino Acids, 20 μM 2-mercaptoethanol) in the presence of BD GolgiPlug™ (1:1000) for 2 hr at 37°C. Cells were then washed, incubated with Fc block, and stained for surface markers PD-1, PSGL-1, and Ly6C as well as LIVE/DEAD™ Fixable Aqua. Cells were next fixed with 2% paraformaldehyde in PBS for 30 min at 4°C, followed by permeabilization with 1X eBioscience™ Permeabilization Buffer for 15 min 4°C. Intracellular staining was performed for CD4, CD44, CXCR5, IFN-γ (XMG1.2), and IL-21 (1 μg of IL-21R-Fc Chimera Protein; R&D Systems, 596-MR-100) in permeabilization buffer for 40 min at 4°C, followed by secondary staining with PE anti-human IgG (Jackson ImmunoResearch, 109-116-098) for 30 min at 4°C.

Samples from SARS-CoV-2-infected mice were fixed with 4% paraformaldehyde for 30 min at room temperature before removal from BSL3 facility. Samples were acquired on a CytoFLEX S (Beckman Coulter) and analyzed using FlowJo software (BD).

### Immunofluorescence

MedLN were fixed with 4% paraformaldehyde in PBS for 4 hr at 4°C, followed by cryopreservation with 20% sucrose in PBS for 2 hr at 4°C. MedLN were snap-frozen in optimal cutting temperature compound and stored at -80°C. Tissues were cut into 7-μm sections and blocked with 10% rat serum in staining solution (PBS with 1% BSA and 0.1% Tween-20) for 1 hr at room temperature. Sections were stained with BV421 anti-PSGL-1 (2PH1) and AF647 anti-CD4 (RM4-5) or BV421 anti-B220 (RA3-6B2) and AF647 anti-GL7 (GL7) along with Fc block, biotinylated anti-mouse IgG2c (5.7), and rabbit anti-T-bet (E4I2K; Cell Signaling Technology, 97135S) overnight at 4°C. Secondary staining was performed with AF594 streptavidin and AF488 goat anti-rabbit IgG for 1 hr at room temperature. Images were acquired on a Nikon TiE Spinning Disk Confocal Microscope with the 30× objective and analyzed with ImageJ.

### Statistical analysis

Data analysis was performed using GraphPad Prism 9 unless otherwise indicated. Data were analyzed using Student’s two-tailed, unpaired t-test; or Welch’s two-tailed, unpaired t-test, as indicated. P < 0.05 was considered statistically significant.

## Results and Discussion

Multiple non-Tfh CD4^+^ T cell populations are induced in lung draining LNs by viral infection Our previous work demonstrated that high-affinity, neutralizing antibodies to SARS-CoV-2 can be produced through Tfh-independent mechanisms, even though certain antibody isotypes (e.g., IgG1) are impaired (11). We therefore analyzed the CD4^+^ T cell populations induced to SARS-CoV-2 at 7 days post-infection (dpi) in mediastinal lymph nodes (medLN) of *Bcl6*^fl/fl^ control and *Bcl6*^fl/fl^*Cd4*^Cre^ Tfh-deficient mice (15). We utilized a previously described model of intratracheal human ACE2 overexpression to enable intranasal SARS-CoV-2 infection of mice (20). While total CD4^+^ T cell counts were unaffected in *Bcl6*^fl/fl^*Cd4*^Cre^ mice, activated CD44^+^CD4^+^ T cell counts were reduced, consistent with the loss of Tfh cells (Fig. 1, A and B). We classified PD-1^lo^CXCR5^lo^CD44^+^CD4^+^ T cells by their expression of PSGL-1 and Ly6C (Fig. 1C). These markers have previously been used in acute LCMV and influenza virus infection to distinguish terminally differentiated PSGL-1^hi^Ly6C^hi^ Th1 cells from a heterogeneous PSGL-1^hi^Ly6C^lo^ Th1 compartment containing memory precursors along with other Th1-related functional subsets (21–23). We therefore defined PSGL-1^hi^Ly6C^hi^ cells as terminally differentiated Th1 cells and PSGL-1^hi^Ly6C^lo^ cells as mixed Th1 cells. We also observed a BCL6-dependent PSGL-1^lo^Ly6C^lo^ population within the PD-1^lo^CXCR5^lo^ gate, which has previously been described as pre-Tfh cells (Fig. 1, D and E) (24). Mixed Th1 and terminally differentiated Th1 cells increased in relative frequency among activated CD4^+^ T cells in *Bcl6*^fl/fl^*Cd4*^Cre^ mice, consistent with the loss of Tfh and pre-Tfh cells; however, their absolute numbers were similar between *Bcl6*^fl/fl^ and *Bcl6*^fl/fl^*Cd4*^Cre^ mice, indicating that BCL6 deficiency did not lead to an aberrant increase in Th1 populations (Fig. 1, F to I).

**Figure 1:**
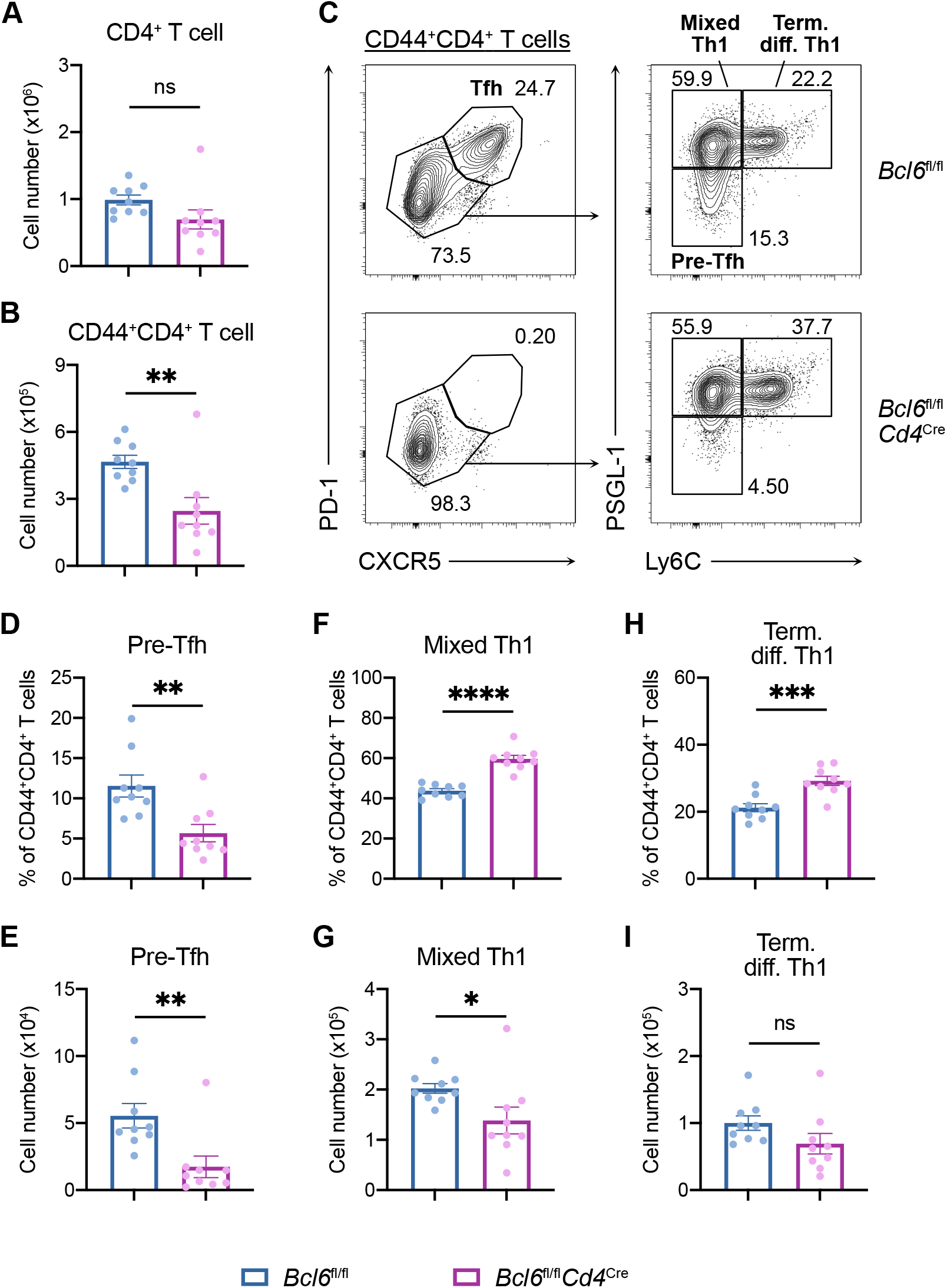
Characterization of CD4^+^ T cell subsets induced by SARS-CoV-2 infection. (**A** to **I**) Flow cytometric analysis of mediastinal lymph nodes (medLN) from SARS-CoV-2-infected mice at 7 dpi. (A and B) Total number of CD4^+^ T cells (**A**) and CD44^+^CD4^+^ T cells (**B**) in *Bcl6*^fl/fl^ (blue) or *Bcl6*^fl/fl^*Cd4*^Cre^ (magenta) mice. (C) Representative gating strategy to define Tfh cells, pre-Tfh cells, mixed Th1 cells, and terminally differentiated Th1 cells from CD44^+^CD4^+^ T cells. (D to I) Frequency among CD44^+^CD4^+^ T cells and total number of pre-Tfh cells (D and E), mixed Th1 cells (F and G), and terminally differentiated Th1 cells (H and I) in *Bcl6*^fl/fl^ (blue) or *Bcl6*^fl/fl^*Cd4*^Cre^ (magenta) mice. Statistical significance was assessed by either two-tailed unpaired t-test or Welch’s t-test, based on the F test for unequal variance. \**P* < 0.05; \*\**P* < 0.01; \*\*\**P* < 0.001; \*\*\*\**P* < 0.0001. ns, not significant. Data are expressed as mean ± SEM. Each symbol represents an individual mouse. Data are aggregated from three independent experiments.

### Th1 cells express costimulatory molecule CD40L

We next evaluated whether these Th1 populations produce CD40L, an effector molecule usually ascribed to Tfh cells (1). While Tfh cells produced the highest levels of CD40L, mixed Th1 cells also produced substantial levels of this effector molecule (Fig. 2, A and B). Tfh cells and mixed Th1 cells comprised the majority of CD40L-expressing CD4^+^ T cells in the medLN of *Bcl6*^fl/fl^ mice, while mixed Th1 cells became the main CD40L-expressing cells in *Bcl6*^fl/fl^*Cd4*^Cre^ mice (Fig. 2C). In *Bcl6*^fl/fl^*Cd4*^Cre^ mice, the frequency of CD40L^+^ cells among CD4^+^ T cells was decreased (Fig. 2D), likely owing to the loss of CD40L-expressing Tfh cells. However, mixed Th1 and terminally differentiated Th1 cells expressed higher levels of CD40L in *Bcl6*^fl/fl^*Cd4*^Cre^ mice compared to *Bcl6*^fl/fl^ mice (Fig. 2, E and F). In the absence of Tfh cells, Th1 subsets may have more opportunities to interact with antigen-presenting cells and experience T cell receptor signaling, which promotes CD40L expression (25). These findings were also generalizable to other models of respiratory viral infection. In *Bcl6*^fl/fl^ mice infected with mouse-adapted influenza virus A/PR/8/34 H1N1 (PR8), mixed Th1 cells also expressed substantial levels of CD40L and constituted a large percentage of CD40L-expressing CD4^+^ T cells (Supplemental Fig. 1, A and B). Therefore, mixed Th1 cells are a significant source of CD40L in the medLN of virally infected *Bcl6*^fl/fl^ and *Bcl6*^fl/fl^*Cd4*^Cre^ mice, suggesting that they may provide contact-dependent help to B cells both in the presence and in the absence of Tfh cells.

**Figure 2:**
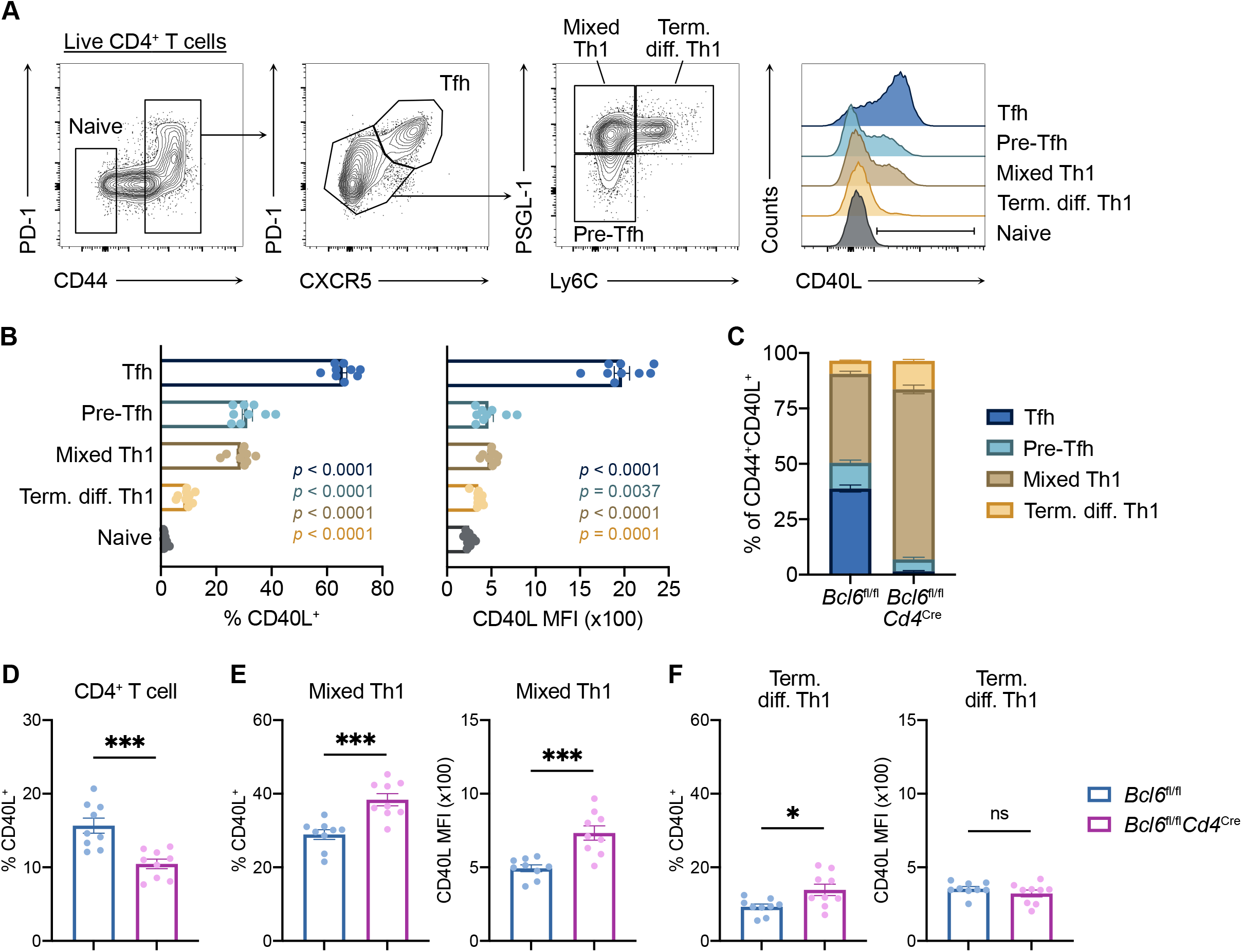
Th1 cells express costimulatory molecule CD40L. (**A** to **C**) Flow cytometric analysis of CD4^+^ T cells in medLN from SARS-CoV-2-infected mice at 7 dpi. (A) Representative gating strategy for quantifying CD40L expression in distinct CD4^+^ T cell subsets. Pre-Tfh cells express a low amount of PD-1 and CXCR5 but are BCL6-dependent (*11*). Mixed Th1 cells classically include Th1 effectors and memory precursors. Terminally differentiated Th1 cells have been shown to express high levels of T-bet, IFN-γ, and GzmB (*8*). (B) Frequency (left) and median fluorescence intensity (MFI) (right) of CD40L expression within CD4^+^ T cell subsets from control *Bcl6*^fl/fl^ mice. Statistical significance was assessed by two-tailed unpaired Welch’s t-test. *P* values for each subset relative to naive CD4^+^ T cells are color-coded. (C) Relative proportions of CD4^+^ T cell subsets among CD44^+^CD40L^+^ cells from *Bcl6*^fl/fl^ or *Bcl6*^fl/fl^*Cd4*^Cre^ mice. (**D** to **F**) Flow cytometric analysis of medLN from *Bcl6*^fl/fl^ (blue) or *Bcl6*^fl/fl^*Cd4*^Cre^ (magenta) mice at 7 dpi with SARS-CoV-2. Frequency of CD44^+^CD40L^+^ cells among CD4^+^ T cells. (E and F) Frequency (left) and MFI (right) of CD40L expression among mixed Th1 cells (E) and terminally differentiated Th1 cells (F). Statistical significance was assessed by either two-tailed unpaired t-test or Welch’s t-test, based on the F test for unequal variance. \**P* < 0.05; \*\*\**P* < 0.001. ns, not significant. Data are expressed as mean ± SEM. Each symbol represents an individual mouse. Data are aggregated from at least two independent experiments.

### Th1 cells express Tfh effector cytokine IL-21

As we had observed that Th1 cells produce Tfh effector molecule CD40L, we next assessed whether they also produce IL-21, a canonical Tfh cytokine that acts at multiple stages to support B cell activation, proliferation, differentiation, and antibody production (1). In SARS-CoV-2-infected *Bcl6*^fl/fl^ mice, Tfh cells produced the highest levels of IL-21 (Fig. 3, A and B). A portion of Tfh cells produced IL-21 together with IFN-γ, which is important for IgG2c class switching (26, 27). Pre-Tfh cells and mixed Th1 cells consisted of IL-21 single producers, IFN-γ single producers, and IL-21/IFN-γ double producers (Fig. 3, C and D). Terminally differentiated Th1 cells were predominantly IFN-γ single producers (Fig. 3E), suggesting that they likely migrate to peripheral tissues to exert their effector function (21). As with CD40L expression, Tfh cells and mixed Th1 cells comprised the majority of IL-21 single-producing and IL-21/IFN-γ double-producing CD4^+^ T cells in *Bcl6*^fl/fl^ mice (Fig. 3, F and G). In *Bcl6*^fl/fl^*Cd4*^Cre^ mice, the frequency of IL-21 single producers decreased, potentially due to the loss of Tfh cells, and mixed Th1 cells became the dominant cytokine-producing population (Fig. 3, F to H). Mixed Th1 and terminally differentiated Th1 cells displayed similar cytokine profiles between *Bcl6*^fl/fl^ and *Bcl6*^fl/fl^*Cd4*^Cre^ mice (Fig. 3, I and J). Thus, mixed Th1 cells express IL-21 in addition to CD40L and are functionally equipped to support an alternative pathway of antibody production to that driven by Tfh cells.

**Figure 3:**
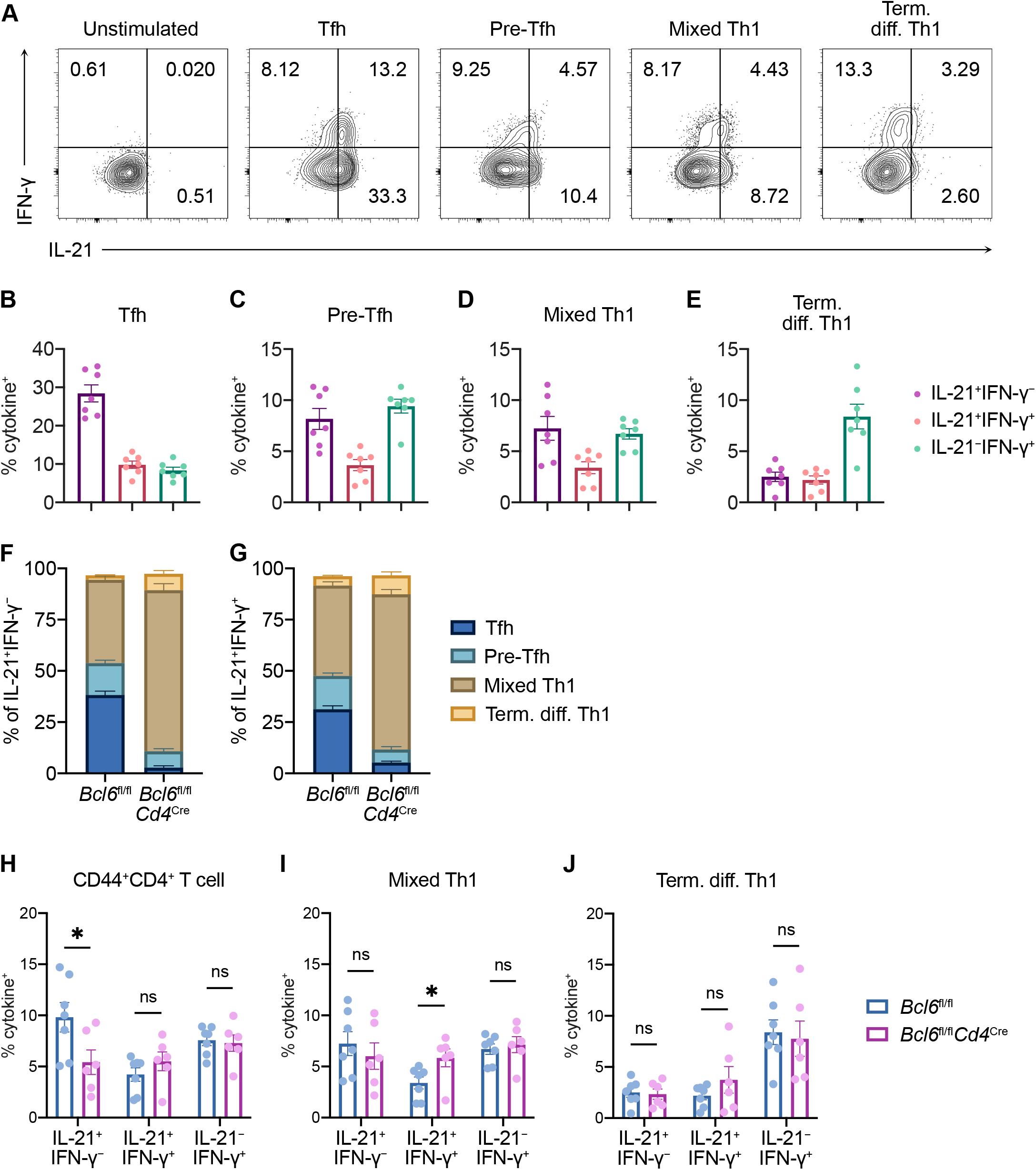
Th1 cells express Tfh effector cytokine IL-21. (**A** to **J**) Flow cytometric analysis of CD4^+^ T cells in medLN from SARS-CoV-2-infected mice at 7 dpi. (A) Representative flow cytometric analysis of IL-21 and IFN-γ protein expression in distinct CD4^+^ T cell subsets from control *Bcl6*^fl/fl^ mice. (B to E) Frequency of IL-21^+^IFN-γ^−^ (purple), IL-21^+^IFN-γ^+^ (red), and IL-21^−^IFN-γ^+^ (teal) expression by intracellular cytokine staining in Tfh cells (B), pre-Tfh cells (C), mixed Th1 cells (D), and terminally differentiated Th1 cells (E) cells from control *Bcl6*^fl/fl^ mice. (F and G) Relative proportions of CD4^+^ T cell subsets among IL-21^+^IFN-γ^−^ (F) and IL-21^+^IFN-γ^+^ (G) cells from *Bcl6*^fl/fl^ or *Bcl6*^fl/fl^*Cd4*^Cre^ mice. (H to J) Frequencies of IL-21^+^IFN-γ^−^, IL-21^+^IFN-γ^+^, and IL-21^−^IFN-γ^+^ expression among CD44^+^CD4^+^ T cells (H), mixed Th1 cells (I) and terminally differentiated Th1 cells (J). Statistical significance was assessed by either two-tailed unpaired t-test or Welch’s t-test, based on the F test for unequal variance. \**P* < 0.05. ns, not significant. Data are expressed as mean ± SEM. Each symbol represents an individual mouse. Data are aggregated from at least two independent experiments.

### Th1 cells co-localize with IgG2c+ B cells following viral infection

We next evaluated whether mixed Th1 cells are sub-anatomically positioned to provide help to B cells. In addition to its role in adhesion, PSGL-1 mediates chemotaxis to CCL21 and CCL19, therefore helping naïve CD4^+^ T cells home to secondary lymphoid organs (28). During their differentiation, Tfh cells downregulate PSGL-1 and upregulate CXCR5 to enable their migration into B cell follicles (24, 29). However, as mixed Th1 cells continue to express PSGL-1 and do not upregulate CXCR5, it is likely that these cells would stay proximal to the T cell zone, rather than into the follicle. Immunofluorescence of medLN following PR8 infection showed that Th1 cells (PSGL-1^+^CD4^+^T-bet^+^) co-localized with IgG2c^+^ B cells at the T cell-B cell (T-B) border (Fig. 4). This was true in both *Bcl6*^fl/fl^ and *Bcl6*^fl/fl^*Cd4*^Cre^ mice, suggesting that Th1 cells promote antibody production in parallel with Tfh cells as well as in their absence.

**Figure 4:**
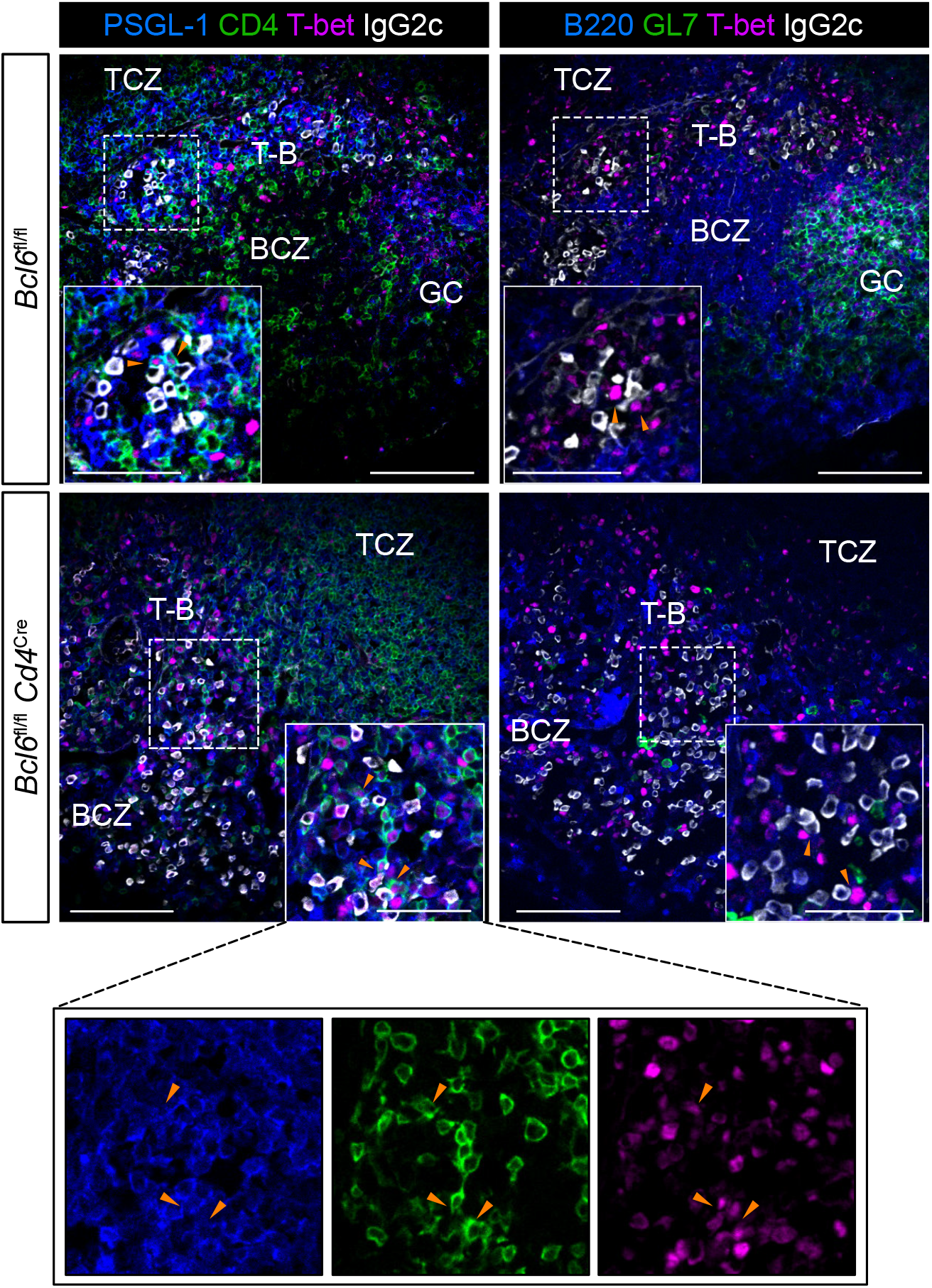
Th1 cells co-localize with IgG2c^+^ B cells following viral infection. Immunofluorescence of serial sections of medLN from *Bcl6*^fl/fl^ (top row) or *Bcl6*^fl/fl^*Cd4*^Cre^ (bottom row) mice infected with PR8 at 14 dpi. Left column: PSGL-1 (blue), CD4 (green), T-bet (magenta), and IgG2c (white). Right column: B220 (blue), GL7 (green), T-bet (magenta), and IgG2c (white). Dashed line demarcates region shown in inset. Orange arrowheads denote Th1 cells interacting with IgG2c^+^ B cells. BCZ, B cell zone. TCZ, T cell zone. T-B, T cell–B cell border. GC, germinal center. Representative images from 3 mice; scale bars, 100 µm and 50 µm (inset).

Taken together, we found that a subset of Th1 cells in LNs expressed CD40L and IL-21 and were positioned at the T-B border to help B cells during viral infection. To distinguish this population from the effector Th1 cells that rapidly leave the LN and function in peripheral tissues (e.g., the lung), we call these cells “LN-Th1” cells. Given our previous work (11) showing that certain isotypes (e.g., IgM, IgG2b, and IgG2c) of spike-specific antibodies could be produced in the absence of GCs following SARS-CoV-2 infection or vaccination, and were capable of virus neutralization, we conclude that these LN-Th1 cells can drive a protective antibody responses to viral infection; our data suggest that this pathway is operational when the GC is impaired, but also when it is intact. We propose that these LN-Th1 cells remain at the T-B border to promote and select antigen-specific B cells, potentially with some limited somatic hypermutation, and thereby act as a parallel pathway to the one within the GC. These findings are consistent with foundational studies highlighting the heterogeneity in T cell responses that drive B cell responses to influenza and more recent work showing an IFN-γ-producing Th1 cell localized to the T-B border during influenza infection and required for anti-viral IgG2c production (14, 30). Although GC-independent responses are sometimes referred to as “extrafollicular” responses, the LN-Th1 pathway we describe here is actually contiguous with the follicle (at the outer border) and therefore we suggest should be distinguished from a classical extrafollicular response (Eisenbarth et al., under revision). An implication of this work is that such a “perifollicular” response can only promote certain antibody isotypes, but these GC-independent antibodies still provide early, effective, and long-lasting viral protection.

## Acknowledgements

We thank all members of the Eisenbarth and Wilen labs for helpful discussions. We would like to acknowledge Benhur Lee, BEI Resources, Joerg Nikolaus, and Yale West Campus Imaging Core for providing critical reagents, resources, and expertise. We thank Yale Environmental Health and Safety for providing necessary training and support for SARS-CoV-2 research. This work was supported by Emergent Ventures Fast Grants (SCE), NIH grant T32GM136651 (JSC, RDC, ES), and NIH grant F30HL149151 (JSC).

## Figure Legends

**Figure S1:**
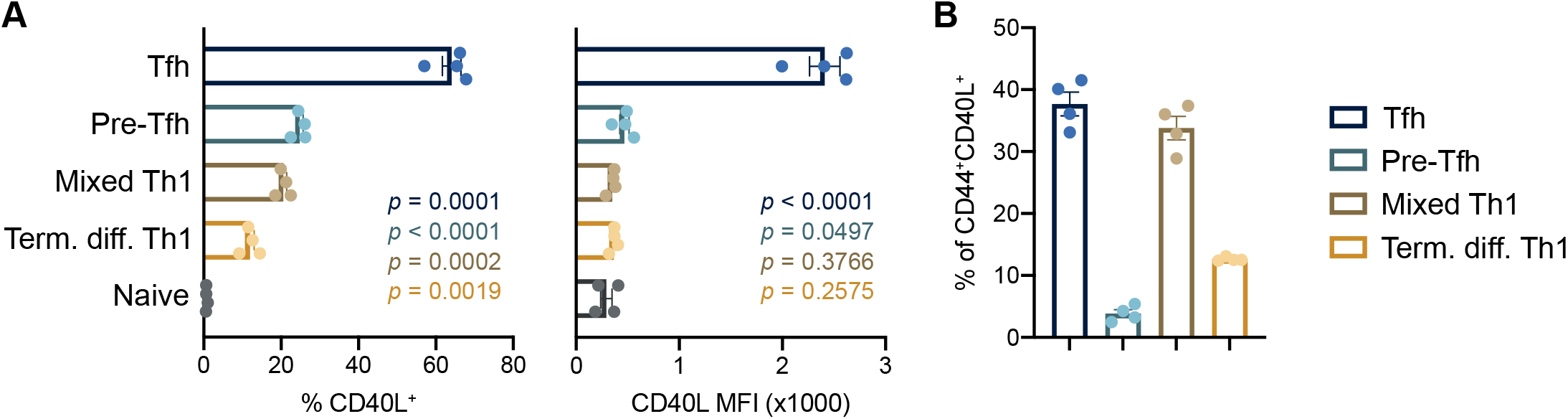
Th1 cells express costimulatory molecule CD40L. (**A** and **B**) Flow cytometric analysis of medLN from *Bcl6*^fl/fl^ mice at 7 dpi with PR8. (D) Frequency (left) and MFI (right) of CD40L expression among CD4^+^ T cell subsets. *P* values for each subset relative to naive CD4^+^ T cells are color-coded. (B) Relative proportions of CD4^+^ T cell subsets among CD44^+^CD40L^+^ cells.

## References

1 Crotty, S. 2019. T Follicular Helper Cell Biology: A Decade of Discovery and Diseases. Immunity 50: 1132–1148.

2 Jacob, J., R. Kassir, and G. Kelsoe. 1991. In situ studies of the primary immune response to (4-hydroxy-3-nitrophenyl)acetyl. I. The architecture and dynamics of responding cell populations. J Exp Med 173: 1165–1175.

3 Liu, Y. J., J. Zhang, P. J. Lane, E. Y. Chan, and I. C. MacLennan. 1991. Sites of specific B cell activation in primary and secondary responses to T cell-dependent and T cell-independent antigens. Eur J Immunol 21: 2951–2962.

4 Weisel, F. J., G. V. Zuccarino-Catania, M. Chikina, and M. J. Shlomchik. 2016. A Temporal Switch in the Germinal Center Determines Differential Output of Memory B and Plasma Cells. Immunity 44: 116–130.

5 Viant, C., T. Wirthmiller, M. A. ElTanbouly, S. T. Chen, E. E. Kara, M. Cipolla, V. Ramos, T. Y. Oliveira, L. Stamatatos, and M. C. Nussenzweig. 2021. Germinal center-dependent and -independent memory B cells produced throughout the immune response. J Exp Med 218: e20202489.

6 Jacob, J., and G. Kelsoe. 1992. In situ studies of the primary immune response to (4-hydroxy-3-nitrophenyl)acetyl. II. A common clonal origin for periarteriolar lymphoid sheath-associated foci and germinal centers. J Exp Med 176: 679–687.

7 Cyster, J. G., and C. C. Goodnow. 1995. Antigen-induced exclusion from follicles and anergy are separate and complementary processes that influence peripheral B cell fate. Immunity 3: 691–701.

8 Toellner, K. M., A. Gulbranson-Judge, D. R. Taylor, D. M. Sze, and I. C. MacLennan. 1996. Immunoglobulin switch transcript production in vivo related to the site and time of antigen-specific B cell activation. J Exp Med 183: 2303–2312.

9 Luther, S. A., A. Gulbranson-Judge, H. Acha-Orbea, and I. C. MacLennan. 1997. Viral superantigen drives extrafollicular and follicular B cell differentiation leading to virus-specific antibody production. J Exp Med 185: 551–562.

10 Chan, T. D., D. Gatto, K. Wood, T. Camidge, A. Basten, and R. Brink. 2009. Antigen affinity controls rapid T-dependent antibody production by driving the expansion rather than the differentiation or extrafollicular migration of early plasmablasts. J Immunol 183: 3139– 3149.

11 Chen, J. S., R. D. Chow, E. Song, T. Mao, B. Israelow, K. Kamath, J. Bozekowski, W. A. Haynes, R. B. Filler, B. L. Menasche, J. Wei, M. M. Alfajaro, W. Song, L. Peng, L. Carter, J. S. Weinstein, U. Gowthaman, S. Chen, J. Craft, J. C. Shon, A. Iwasaki, C. B. Wilen, and S. C. Eisenbarth. 2021. High-affinity, neutralizing antibodies to SARS-CoV-2 can be made without T follicular helper cells. Science Immunology 7.

12 Zhang, B., E. Liu, J. A. Gertie, J. Joseph, L. Xu, E. Y. Pinker, D. A. Waizman, J. Catanzaro, K. H. Hamza, K. Lahl, U. Gowthaman, and S. C. Eisenbarth. 2020. Divergent T follicular helper cell requirement for IgA and IgE production to peanut during allergic sensitization. Science Immunology 5.

13 Miyauchi, K., A. Sugimoto-Ishige, Y. Harada, Y. Adachi, Y. Usami, T. Kaji, K. Inoue, H. Hasegawa, T. Watanabe, A. Hijikata, S. Fukuyama, T. Maemura, M. Okada-Hatakeyama, O. Ohara, Y. Kawaoka, Y. Takahashi, T. Takemori, and M. Kubo. 2016. Protective neutralizing influenza antibody response in the absence of T follicular helper cells. Nat Immunol 17: 1447–1458.

14 Mendoza, A., W. T. Yewdell, B. Hoyos, M. Schizas, R. Bou-Puerto, A. J. Michaels, C. C. Brown, J. Chaudhuri, and A. Y. Rudensky. 2021. Assembly of a spatial circuit of T-bet-expressing T and B lymphocytes is required for antiviral humoral immunity. Sci Immunol 6: eabi4710.

15 Hollister, K., S. Kusam, H. Wu, N. Clegg, A. Mondal, D. V. Sawant, and A. L. Dent. 2013. Insights into the Role of Bcl6 in Follicular Th Cells Using a New Conditional Mutant Mouse Model. The Journal of Immunology 191: 3705–3711.

16 Lee, P. P., D. R. Fitzpatrick, C. Beard, H. K. Jessup, S. Lehar, K. W. Makar, M. Pérez-Melgosa, M. T. Sweetser, M. S. Schlissel, S. Nguyen, S. R. Cherry, J. H. Tsai, S. M. Tucker, W. M. Weaver, A. Kelso, R. Jaenisch, and C. B. Wilson. 2001. A Critical Role for Dnmt1 and DNA Methylation in T Cell Development, Function, and Survival. Immunity 15: 763–774.

17 Wei, J., M. M. Alfajaro, P. C. DeWeirdt, R. E. Hanna, W. J. Lu-Culligan, W. L. Cai, M. S. Strine, S.-M. Zhang, V. R. Graziano, C. O. Schmitz, J. S. Chen, M. C. Mankowski, R. B. Filler, N. G. Ravindra, V. Gasque, F. J. de Miguel, A. Patil, H. Chen, K. Y. Oguntuyo, L. Abriola, Y. V. Surovtseva, R. C. Orchard, B. Lee, B. D. Lindenbach, K. Politi, D. van Dijk, C. Kadoch, M. D. Simon, Q. Yan, J. G. Doench, and C. B. Wilen. 2021. Genome-wide CRISPR Screens Reveal Host Factors Critical for SARS-CoV-2 Infection. Cell 184: 76-91.e13.

18 Pillai, P. S., R. D. Molony, K. Martinod, H. Dong, I. K. Pang, M. C. Tal, A. G. Solis, P. Bielecki, S. Mohanty, M. Trentalange, R. J. Homer, R. A. Flavell, D. D. Wagner, R. R. Montgomery, A. C. Shaw, P. Staeheli, and A. Iwasaki. 2016. Mx1 reveals innate pathways to antiviral resistance and lethal influenza disease. Science 352: 463–466.

19 Koguchi, Y., T. J. Thauland, M. K. Slifka, and D. C. Parker. 2007. Preformed CD40 ligand exists in secretory lysosomes in effector and memory CD4+ T cells and is quickly expressed on the cell surface in an antigen-specific manner. Blood 110: 2520–2527.

20 Israelow, B., E. Song, T. Mao, P. Lu, A. Meir, F. Liu, M. M. Alfajaro, J. Wei, H. Dong, R. J. Homer, A. Ring, C. B. Wilen, and A. Iwasaki. 2020. Mouse model of SARS-CoV-2 reveals inflammatory role of type I interferon signaling. Journal of Experimental Medicine 217: e20201241.

21 Marshall, H. D., A. Chandele, Y. W. Jung, H. Meng, A. C. Poholek, I. A. Parish, R. Rutishauser, W. Cui, S. H. Kleinstein, J. Craft, and S. M. Kaech. 2011. Differential Expression of Ly6C and T-bet Distinguish Effector and Memory Th1 CD4+ Cell Properties during Viral Infection. Immunity 35: 633–646.

22 Marshall, H. D., J. P. Ray, B. J. Laidlaw, N. Zhang, D. Gawande, M. M. Staron, J. Craft, and S. M. Kaech. The transforming growth factor beta signaling pathway is critical for the formation of CD4 T follicular helper cells and isotype-switched antibody responses in the lung mucosa. eLife 4: e04851.

23 Pepper, M., A. J. Pagán, B.Z. Igyártó, J. J. Taylor, and M. K. Jenkins. 2011. Opposing Signals from the Bcl6 Transcription Factor and the Interleukin-2 Receptor Generate T Helper 1 Central and Effector Memory Cells. Immunity 35: 583–595.

24 Poholek, A. C., K. Hansen, S. G. Hernandez, D. Eto, A. Chandele, J. S. Weinstein, X. Dong, J. M. Odegard, S. M. Kaech, A. L. Dent, S. Crotty, and J. Craft. 2010. In Vivo Regulation of Bcl6 and T Follicular Helper Cell Development. The Journal of Immunology 185: 313– 326.

25 Jaiswal, A. I., C. Dubey, S. L. Swain, and M. Croft. 1996. Regulation of CD40 ligand expression on naive CD4 T cells: a role for TCR but not co-stimulatory signals. Int Immunol 8: 275–285.

26 Snapper, C. M., and W. E. Paul. 1987. Interferon-gamma and B cell stimulatory factor-1 reciprocally regulate Ig isotype production. Science 236: 944–947.

27 Reinhardt, R. L., H.-E. Liang, and R. M. Locksley. 2009. Cytokine-secreting follicular T cells shape the antibody repertoire. Nat Immunol 10: 385–393.

28 Veerman, K. M., M. J. Williams, K. Uchimura, M. S. Singer, J. S. Merzaban, S. Naus, D. A. Carlow, P. Owen, J. Rivera-Nieves, S. D. Rosen, and H. J. Ziltener. 2007. Interaction of the selectin ligand PSGL-1 with chemokines CCL21 and CCL19 facilitates efficient homing of T cells to secondary lymphoid organs. Nat Immunol 8: 532–539.

29 Haynes, N. M., C. D. C. Allen, R. Lesley, K. M. Ansel, N. Killeen, and J. G. Cyster. 2007. Role of CXCR5 and CCR7 in Follicular Th Cell Positioning and Appearance of a Programmed Cell Death Gene-1High Germinal Center-Associated Subpopulation. The Journal of Immunology 179: 5099–5108.

30 Elsner, R. A., D. N. Ernst, and N. Baumgarth. 2012. Single and coexpression of CXCR4 and CXCR5 identifies CD4 T helper cells in distinct lymph node niches during influenza virus infection. J Virol 86: 7146–7157.

